# Ectopically Expressed Rhodopsin is Not Sensitive to X-rays

**DOI:** 10.1101/2023.08.03.551542

**Authors:** Kelli Cannon, Aundrea Bartley, Lynn Dobrunz, Mark Bolding

**Affiliations:** Department of Vision Sciences, School of Optometry, University of Alabama at Birmingham, Birmingham, AL, United States; Department of Neurobiology, Heersink School of Medicine, University of Alabama at Birmingham, Birmingham, AL, United States; Department of Radiology, Heersink School of Medicine, University of Alabama at Birmingham, Birmingham, AL, United States

**Keywords:** Rhodopsin, X-rays, X-genetics, optogenetics, noninvasive neuromodulation

## Abstract

Visual perception of X-radiation is a well-documented, but poorly understood phenomenon. Early literature implicates scotopic rod cells and rod opsin in X-ray detection, however, evidence suggests that X-rays excite the retina via a different mechanism than visible light. While rhodopsin’s role in X-ray perception is unclear, the possibility that it could function as an X-ray receptor has led to speculation that it could act as a transgenically expressed X-ray receptor. If so, it could be used to transduce transcranial X-ray signals and control the activity of genetically targeted populations of neurons in a less invasive version of optogenetics, X-genetics. Here we investigate whether human rhodopsin (hRho) is capable of transducing X-ray signals when expressed outside of the retinal environment. We use a live-cell cAMP GloSensor luminescence assay to measure cAMP decreases in hRho-expressing HEK293 cells in response to visible light and X-ray stimulation. We show that cAMP GloSensor luminescence decreases are not observed in hRho-expressing HEK293 cells in response to X-ray stimulation, despite the presence of robust responses to visible light. Additionally, irradiation had no significant effect on cAMP GloSensor responses to subsequent visible light stimulation. These results indicate that ectopically expressed rhodopsin does not function as an X-ray receptor, and suggest that it is not capable of transducing transcranial X-ray signals into neural activity for X-ray mediated, genetically targeted neuromodulation.

## Introduction

Neuromodulation has entered an exciting new era with the emergence of optogenetics and other genetically targeted technologies, enabling researchers to manipulate neural activity in a cell-type-specific manner. Optogenetics utilizes visible light to activate photoreceptors expressed in targeted neural populations. Despite its proven utility, the use of visible light stimulation presents certain drawbacks, primarily due to its limited penetration and high scattering within tissue. The standard approach for in vivo optogenetics experiments involves invasive transcranial implantation of an optic fiber to deliver light to the desired brain region. The stimulation is confined to a narrow area a few millimeters wide surrounding the fiber tip, as substantial scattering of light hampers its efficient delivery, leading to excessive tissue warming when higher light levels are used Deng et al. (2014); Foutz et al. (2012). This spatial restriction also necessitates the implantation of multiple fibers when targeting multiple brain areas. As such, the use of visible light imposes significant limitations on optogenetics studies.

Given the shortcomings of visible light for neural stimulation, researchers have begun investigating the use of alternative forms of stimulation, including X-rays, for genetically targeted neuromodulation Berry et al. (2015). X-rays are well suited for transmission through tissue, including skull, eliminating the need for transcranial implants for stimulation delivery. X-rays are also substantially less scattered by tissue Matsubara et al. (2021), allowing for efficient focusing of stimulation to target deep brain structures, even in large animals, while minimizing off-target stimulation of intervening tissue. Multiple brain areas can easily be targeted by irradiating the whole brain with an unfocused X-ray beam or by utilizing multiple, focused X-ray sources. These advantages promote X-radiation as potentially a superior form of stimulation for genetically targeted neural control.

The use of X-rays for genetically targeted neural stimulation requires a means to detect X-rays using a genetically targeted receptor. To address this, certain research groups have investigated the utilization of radioluminescent materials, which emit visible light when excited by X-rays. These materials, when combined with opsins, have shown promise in achieving “X-ray optogenetic” control of neural activity Bartley et al. (2021); Chen et al. (2021); Matsubara et al. (2021). However, the light output of these materials when excited by biologically tolerable levels of X-rays is very low compared to the light levels delivered by fiber optic implants, and it is not clear whether radioluminescent materials can be used noninvasively for X-ray optogenetic neuromodulation. Alternatively, a receptor directly sensitive to X-rays or their biochemical byproducts would eliminate the requirement for visible light and possibly boost X-ray transduction efficiency to allow effective control of neural activity using lower X-ray doses. Our group has discovered that LITE-1, an unusual UV photoreceptor found in C. elegans, can function as an X-ray receptor, mediating “X-genetic” control of muscle cells ectopically expressing LITE-1 Cannon et al. (2023). These initial X-ray optogenetics and X-genetics studies offer proof-of-concept for the application of X-rays in genetically targeted neuromodulation, however further efforts are necessary to optimize and refine these techniques to include a variety of receptors to provide different response properties and ensure their widespread utility.

Another photoreceptor, mammalian rhodopsin, has historically been implicated in visual responses to X-rays, including X-ray phosphenes and ERG responses in humans and a variety of animal models Brandes and Dorn (1897); Dawson and Smith (1959); Kimeldorf and Hunt (1965); Lipetz (1955); Newell and Borley (1941). The ubiquitous finding that the retina must be in a scotopic, or dark-adapted, state in order to detect X-rays led to speculation that scotopically active rod opsin—opposed to photopically active cone opsins—functions as the mediating receptor Bachofer and Wittry (1961, 1962); Kimeldorf and Hunt (1965); Lipetz (1955). However, some lines of evidence, including the finding that X-rays do not bleach rod opsin at the intensities used to elicit perceptual and ERG responses Baldwin et al. (1963); Doly et al. (1980), suggest that different mechanisms underlie retinal detection of X-rays, compared to visible light. One recent study revisited the X-ray ERG and stipulated that rhodopsin might be a suitable receptor for X-genetic control of neural activity Getzin et al. (2017). No study to date, however, has directly tested whether rhodopsin is capable of transducing X-ray signals when expressed outside of the specialized environment of the retina. The present study tests whether human rhodopsin is activated by X-rays by measuring G-protein mediated cAMP GloSensor luminescence responses induced by visible light and X-ray stimulation in human embryonic kidney (HEK293) cells expressing hRho. We show that robust hRho-mediated decreases in cAMP levels are observed in response to visible light, but no responses are observed in response to X-rays. This suggests that visual responses to X-rays may not be mediated by rhodopsin, despite long-standing speculation, indicating that rhodopsin is unlikely to be usable as an X-ray receptor for X-genetics.

## Methods

Human embryonic kidney cells (HEK293) were obtained from the lab of Dr. Heinrich Matthies at the University of Alabama at Birmingham. pcDNA3 Rod Opsin plasmid (hRho) was obtained from Addgene (plasmid #109361). GloSensor-22F plasmid was obtained from Promega Corporation (E2301). Cells were maintained in DMEM with 10% fetal bovine serum (FBS), 6 mM L-glutamine (LG), and 1% penicillin/streptomycin at 37°C and 5% CO2.

### Immunolabeling for detection of hRho

#### Cell Preparation and Transfection

HEK293 cells were plated in T25 flasks in DMEM with 10% FBS and 6 mM LG the day before transfection and transfected with pcDNA3 Rod Opsin plasmid DNA using Lipofectamine 2000, according to manufacturer’s instructions. Approximately 6 hrs after transfection under dim red light, cells were resuspended in fresh antibiotic-free DMEM with 10% FBS, 6 mM LG, and 10 µM 9-cis-retinal and replated on on poly-d-lysine coated glass coverslips in a 12 well plate at a density of 150,00 cells per coverslip for immunofluorescence assay.

#### Immunofluorescence Assay

Immunostaining for 1D4 was used to detect expression of hRho after transfection. Approximately 24 hrs after transfection, coverslips were rinsed once with 1x Dulbecco’s phosphate buffered saline with calcium and magnesium (DPBS), fixed with 4% paraformaldehyde, then rinsed 3 more times with DPBS before storage in DPBS at 4°C. Immunolabeling of coverslips was performed using 10% normal goat serum blocking buffer with 0.2% Triton-X100, mouse 1D4 monoclonal antibody (obtained from the lab of Dr. Alecia Gross at the University of Alabama at Birmingham) at 1:1000 dilution, rinsing 3 times with DPBS, then applying AlexaFluor568 goat anti-mouse secondary antibody (Life Technologies A11001) at a 1:1000 dilution. After 3 further DPBS rinses, cell nuclei were stained with 1 µg/mL DAPI. After a final DPBS rinse, coverslips were mounted on glass slides. Fluorescence images of slides were obtained on an ECHO Revolution microscope.

### Live-Cell GloSensor Assays

#### Cell Preparation and Transfection

HEK293 cells were plated in T25 flasks in DMEM with 10% FBS and 6 mM LG the day before transfection and transfected with pcDNA3 Rod Opsin and pGloSensor-22F plasmids in a 1:1 ratio using Lipofectamine 2000. Control cells were transfected with pGloSensor-22F only. Approximately 6 hrs after transfection under dim red light, cells were resuspended in fresh antibiotic-free DMEM with 10% FBS, 6 mM LG, and 10 µM 9-cis-retinal and replated in white flat-bottom 96-well plates, with 3-4 wells per condition for technical replicates. The next day under dim red light, cells in 96 well plate were equilibrated in phenol-red-free L-15 medium with 1% FBS, 6 mM LG, 10 µM 9-cis-retinal, and 2 mM d-luciferin outside of the incubator for 1 - 1.5 hrs before conducting the GloSensor assay.

#### Stimulation

Four stimulation conditions were used: 1) negative control cells unexposed to either X-rays or visible light, 2) positive control cells exposed to visible light only, 3) cells exposed to moderate dose rate X-rays, and 4) cells exposed to high dose rate X-rays. For visible light stimulation, approximately 10^15^ photons/mm^2^ visible light was delivered to cells using a 1.6 s pulse of 0.27 mW/mm^2^ 470 nm light from a custom LED array. For the high dose rate X-ray stimulation, 0.34 Gy X-radiation was delivered to cells using a 15 s pulse of 1.36 Gy/min X-ray stimulation from an enclosed X-ray unit (X-RAD 320, W target, no filters) operated at 150 kV/12.5 mA. The dose rate at the level of the plate in the X-RAD 320 was measured using a UNIDOS E dosimeter (PTW-Frieburg, model T10010). For the low dose rate X-ray stimulation, 0.12 mm Pb equivalent acrylic was placed over the designated wells to deliver 0.08 Gy X-radiation at a rate of 0.32 Gy/min. The half of the 96 well plate designated for the unexposed and visible light conditions was placed inside a 0.5 mm Pb equivalent glove, which was verified to reduce dose rate to 0.00 Gy/min.

#### Luminescence Recordings

Luminescence was measured using a plate reader (SpectraMax M3) set to take a 1 s exposure of each well every 90 s. Five baseline readings were collected, then the plate was ejected and 2 µM forskolin was added to cells in order to artificially elevate baseline cAMP to facilitate detection of a stimulation-elicited decrease in cAMP level. Cells were returned to the plate reader, and luminescence measurements were collected, as before. Once luminescence signals were observed to stabilize, the plate was ejected and positive control cells were exposed to visible light stimulation, then experimental cells were exposed to X-ray stimulation. After stimulation, the cells were returned to the plate reader to record luminescence for 11 additional cycles.

Irradiated and unexposed control cells were then exposed to visible light stimulation before returning to the plate reader for 11 more cycles. The assay was conducted six times using cells of different passage numbers.

#### Data Analysis

Data analysis was performed using custom Matlab (2021b) code. GloSensor luminescence data was normalized to baseline, set as the luminescence measurement collected immediately prior to the administration of stimulation. Responses were calculated as the largest deviation from baseline over the 11 measurements collected after stimulation. For statistical analysis of the data from the visible and X-ray stimulation experiment, a factorial ANOVA was performed, followed by a series of two-tailed unpaired t tests to compare mean responses to unexposed controls and to hRho-lacking controls. Bonferroni correction was used to adjust p-values for multiple comparisons. The data from the post-irradiation photoresponse assay was found to be heteroscedastic. Therefore a nonparametric Scheirer-Ray-Hare test was used, followed by a series of two-tailed unpaired Mann Whitney U tests.

## Results

In order to test rhodopsin’s utility for X-genetics, this study used HEK293 cells, which are known to express the downstream signaling components needed to detect rhodopsin photoresponses Atwood et al. (2011). HEK293 cells were transfected with pcDNA3 Rod Opsin, and hRho was allowed to express for 24 hrs. Immunostaining using 1D4 antibody, which targets the C-terminus of hRho, revealed a high transfection efficiency and robust expression of hRho (Figure 1). As a Gi/o/t-coupled GPCR, hRho has been shown to reliably produce visible light-triggered decreases in cAMP levels when expressed in HEK293 cells Ballister et al. (2018). To measure hRho activation in response to stimulation, we performed a live-cell GloSensor cAMP luminescence assay on HEK293 cells transfected with hRho and control cells lacking hRho. As expected, we found that a 10^15^ photons/mm^2^ pulse of visible (470 nm) light reduced GloSensor cAMP luminescence signals of hRho expressing HEK293 cells (0.235±0.056) relative to unexposed control cells (0.838±0.160), a response not observed in cells lacking hRho (light exposed 0.672±0.123; unexposed 0.847±0.202; Figure 2). There was a significant interaction effect of hRho expression and stimulation condition on cAMP luminescence responses, F(40) = 8.18, p = 2.31e-4. Only the responses of visible light exposed hRho expressing cells were significantly different from unexposed (t(6) = 8.70, p = 1.07e-4) and hRho lacking controls (t(7) = 7.91, p = 9.96e-5).

**Fig. 1.**
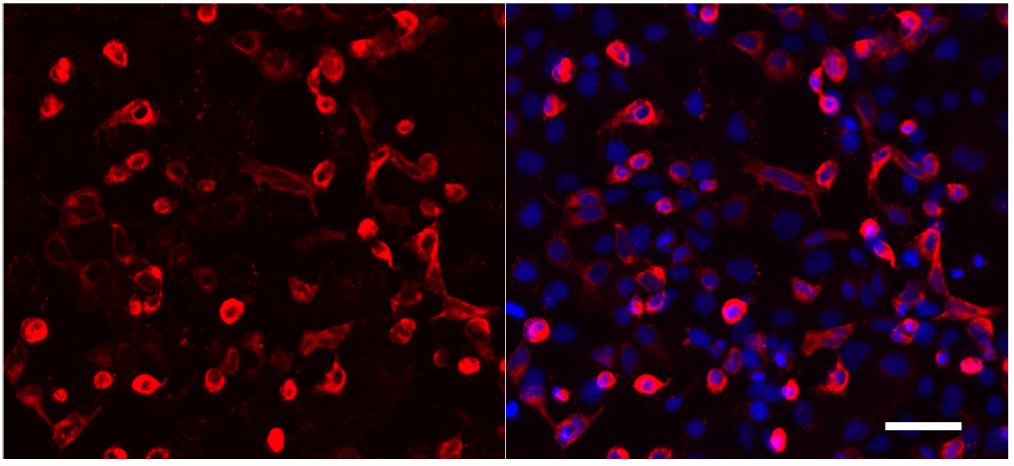
HEK293 cells robustly express hRho. Cells were fixed 24 hrs after transfection with hRho and labeled with 1D4 monoclonal antibody and AlexaFluor568 secondary antibody (red). Nuclei were stained with DAPI (blue). Scale bar indicates 100 µm.

**Fig. 2.**
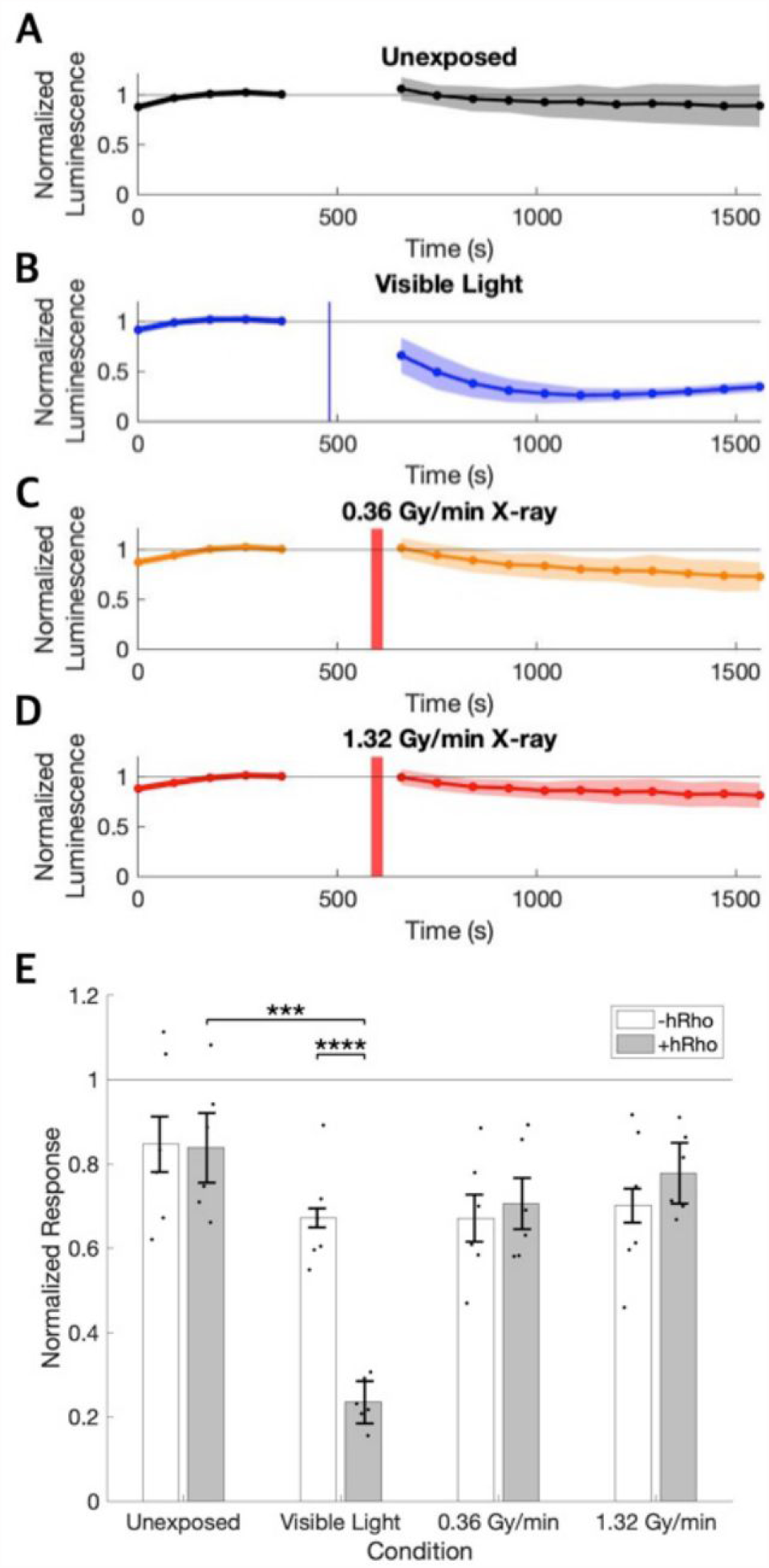
hRho expressed in HEK293 cells responds to visible light stimulation, but not X-rays. A-D) Luminescence signals normalized to baseline and averaged across technical and biological replicates are shown before and after stimulation for hRho expressing cells unexposed to either type of stimulation (black), cells exposed to visible light (470 nm) stimulation, and cells exposed to 0.28 Gy/min (orange) and 1.0 Gy/min (red) X-ray stimulation. Vertical blue and red lines indicate approximate timing of visible light and X-ray pulses, respectively. Shading indicates standard error of the mean traces. C) The peak change in luminescence signal after stimulation normalized to the pre-stimulation baseline signal is shown for cells with and without hRho for each stimulation condition. Dots show the mean responses across technical replicates for a single experiment. Error bars show standard error. *** indicates p < 0.001, **** indicates p < 0.0001. N = 6 for each condition..

To determine whether hRho expressing HEK293 cells could be activated by X-ray stimulation, GloSensor cAMP luminescence responses were recorded for cells exposed to two levels of X-ray stimulation, 0.32 Gy/s and 1.36 Gy/s. No responsiveness to 0.32 or 1.36 Gy/min X-ray stimulation was observed in hRho expressing cells (0.32 Gy/min 0.706±0.138; 1.36 Gy/min 0.778±0.099) compared to hRho lacking cells (0.32 Gy/min 0.671±0.149; 1.36 Gy/min 0.701±0.177; Figure 2), providing strong evidence that hRho is not capable of transducing X-ray signals when expressed outside of the retina. Additionally, no significant difference was observed between hRho expressing cells exposed to 0.32 or 1.36 Gy/min X-rays compared to hRho expressing cells unexposed to light or X-ray stimulation (0.32 Gy/min t(10) = -1.531, p = 0.157; 1.36 Gy/min t(8) = -0.777, p = 0.459). These data suggest that X-rays are not an effective activator of ectopically expressed hRho.

To determine whether X-irradiation had an effect on hRho photoresponses, cells exposed to X-radiation were subsequently exposed to a pulse of 10^15^ photons/mm^2^ visible light. Light stimulation elicited a decrease in GloSensor cAMP luminescence (0.274±0.064) relative to dark exposed controls (0.904±0.033; Figure 3), verifying that hRho was still functional and unbleached in X-ray exposed cells. This difference in responses was significant, U = 16, p = 0.029. The mean response amplitude of irradiated cells to visible light was not significantly different from that of unirradiated cells (0.298±0.046; U = 7, p = 0.886), demonstrating that irradiation had no effect on photoresponses.

**Fig. 3.**
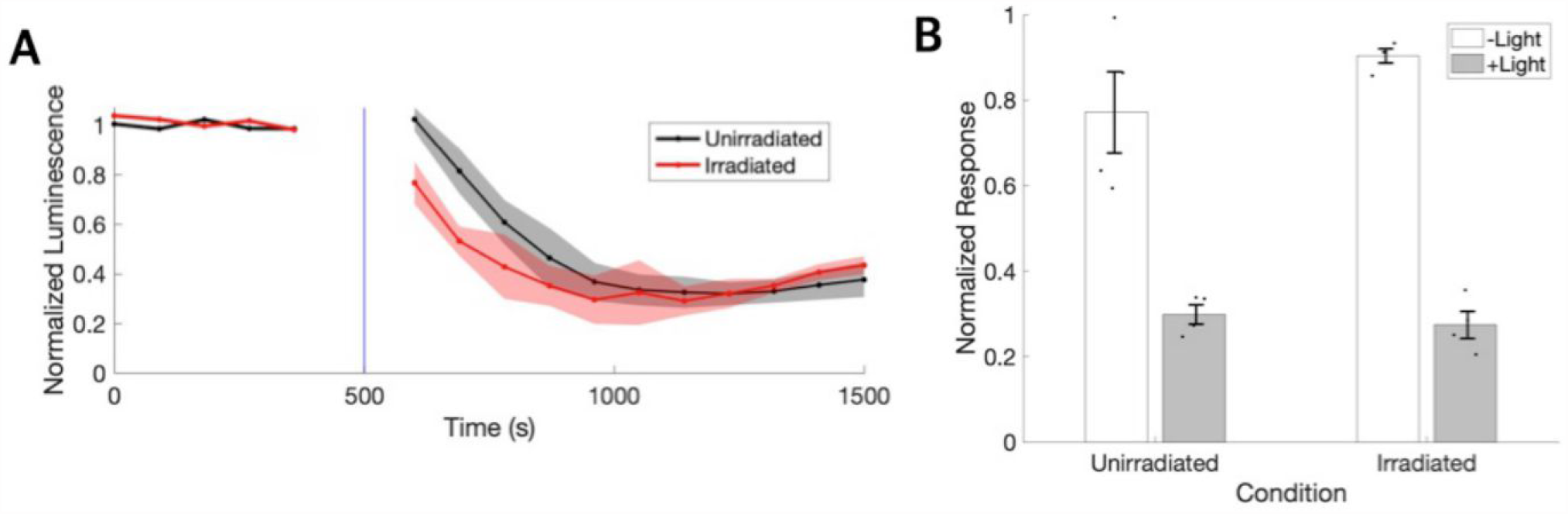
Irradiated cells have an unaltered response to visible light. A) Luminescence signals normalized to baseline and averaged across technical and biological replicates are shown before and after stimulation for hRho expressing cells exposed to 1015 photons/mm2 visible (470 nm) light. Vertical blue line indicates the approximate timing of the visible light pulse. Shading indicates standard error of the mean traces. B) The peak deviation of normalized luminescence from the pre-stimulation baseline value is averaged across replicates for previously irradiated and unirradiated cells exposed to a pulse of visible light, compared to dark exposed controls. Dots represent values for biological replicates. Error bars indicate standard error. N = 4 for each condition.

## Discussion

This study shows that human rod opsin, when expressed outside of the specialized system of the retina, is not responsive to 0.32 or 1.36 Gy/min X-ray stimulation, despite retaining its sensitivity to visible light. We observed a 76% decrease in GloSensor cAMP luminescence in response to 10^15^ photons/mm^2^ visible light, comparable to the 55% decrease observed by Ballister et al. under similar conditions Ballister et al. (2018). Additionally, we found that visual response amplitudes were unaffected by prior irradiation, in line with previous ERG and photochemistry studies showing that X-rays do not bleach rhodopsin Baldwin et al. (1963); Doly et al. (1980).

Investigations of retinal responses to X-rays have employed varying stimulation parameters. X-ray tube voltages vary widely, but voltages as low as 40 kV have been used to elicit responses Getzin et al. (2017); Kimeldorf and Hunt (1965), consistent with the 150 kV tube voltage used in the current study. Early human studies found threshold dose rates for X-ray detection in normal subjects to be much lower than those used here. One study reported threshold dose rates between 0.5 - 1.4 r/min Newell and Borley (1941), and another reported thresholds of 1.6 to 8.7 mr/sec Bornschein et al. (1953). Converted to Gy/min, these dose rates of 4-12 mGy/min or 1-5 mGy/min fall well below the 0.32 and 1.36 Gy/min dose rates used here. Therefore, the X-ray stimulation parameters used in the present study are consistent with those used to elicit X-ray phosphene and ERG responses in retinal studies.

Prior research regarding the effect of retinal irradiation on subsequent visible light responses has yielded mixed results. Some researchers have reported reduced retinal sensitivity to visible light after X-ray exposure in the form of diminished photic ERG b-wave amplitudes and increased thresholds for visible light Baldwin et al. (1963); Lipetz (1955). Others have found sensitivity to visible light to be enhanced after irradiation, reporting decreased visible light thresholds Dawson and Smith (1959); Dawson and Wiederwohl (1965). We found no significant difference in visible light response amplitudes after X-ray stimulation, though it is possible that differences were present at a shorter time point after irradiation than was tested here (15 min).

While this study is not intended to determine the molecular mechanisms underlying visual X-ray perception, it does provide important relevant information, namely that either 1) the specialized system of the retina (e.g., the extremely high density of hRho) is needed for hRho to transduce X-ray signals, or 2) a different molecular receptor operating downstream from hRho in the rod phototransduction pathway is responsible for transducing X-ray signals in the retina.

Rhodopsin expression in the retina is in a highly specialized arrangement with exceptionally high expression densities in the stacks of membrane discs in rod outer segments. This arrangement evolved to maximize light detection, allowing for the detection of sparse photons in the dark-adapted state, but is not recapitulated when Rho is expressed ectopically. While hRho showed robust immunolabeling in HEK293 cells, the number and density of hRho in HEK293 cells is still significantly diminished relative to that seen in rod outer segments. Therefore, it may be that hRho is being activated by X-rays in the retina, but responses are only detectable due to hRho’s high density in rod outer segments, which increases the probability of photon detection. Alternatively, Narici et al. proposed a model for a generation method of radiation-induced phosphenes where radiogenic radicals oxidize lipids, generating chemiluminescent photons that go on to activate rhodopsin Narici et al. (2009). Perhaps the high density of lipids or the particular lipid composition of the membrane discs of rod outer segments is needed for rhodopsin to respond to X-rays.

Alternatively, it may be that retinal X-ray detection is not dependent on rhodopsin at all, with X-rays impinging on some other cellular or molecular target involved in scotopic retinal pathways to elicit retinal responses. Previous findings that X-rays do not bleach hRho at the doses used to elicit phosphene and ERG responses Baldwin et al. (1963); Doly et al. (1980) and that X-ray and light ERGs have distinct dynamics Savchenko (1993) are consistent with the explanation that hRho is not involved in retinal X-ray detection. It is possible that molecular targets act downstream of hRho, including those in downstream neurons like rod bipolar cells. For example, TRPM2 is an ion channel that functions in rod ON bipolar cells and some of its isoforms have been found to respond to X-ray stimulation Klumpp et al. (2016) and reactive oxygen species Perraud et al. (2005), a major by-product of irradiation. However, the kinetics of the TRPM2 response to these stimuli are much slower than the kinetics of typical photoreceptor proteins Sumoza-Toledo and Penner (2011). Others have suggested that X-rays or their by-products could be piercing the membranes of rod photoreceptors Savchenko (1993), though it is unclear why this effect would not be seen in cone photoreceptors if this were the case. The possibilities mentioned here are merely speculations, and future molecular studies are needed to elucidate the actual mechanism(s) involved in retinal X-ray detection.

In summary, this study investigated the potential of human rod opsin to transduce X-ray signals when expressed outside of the specialized environment of the retina. Through live-cell GloSensor cAMP luminescence assays, we found that hRho-expressing cells did not exhibit any significant response to X-ray stimulation above that seen by hRho-lacking cells, while still displaying robust sensitivity to visible light. This suggests that hRho may not be capable of detecting X-rays, at least not when expressed in non-retinal cells. The results imply that either the specialized retinal environment or a different molecular receptor downstream from hRho is responsible for X-ray perception in the retina. Further research is needed to unravel the molecular mechanisms underlying visual X-ray perception and identify the specific players involved in transducing X-ray signals in the retina. However, this study shows that rhodopsin alone is unlikely to be able to function as an X-ray detector for X-genetic stimulation.

## ACKNOWLEDGEMENTS

HEK293 cells used in this study were generously provided by Dr. Heinrich Matthies. Mouse 1D4 monoclonal antibody was generously provided by Dr. Alecia Gross. The X-RAD 320 X-ray unit was purchased using a Research Facility Improvement Grant, 1 G20RR022807-01, from the National Center for Research Resources, National Institutes of Health.

